# Sleep deprivation selectively up-regulates an amygdala-hypothalamic circuit involved in food reward

**DOI:** 10.1101/245845

**Authors:** Julia S. Rihm, Mareike M. Menz, Heidrun Schultz, Luca Bruder, Leonhard Schilbach, Sebastian M. Schmid, Jan Peters

## Abstract

Sleep loss is associated with increased obesity risk, as demonstrated by correlations between sleep duration and change in body mass index or body fat percentage. Whereas previous studies linked this weight gain to disturbed endocrine parameters after sleep deprivation (SD) or restriction, neuroimaging studies revealed up-regulated neural processing of food rewards after sleep loss in reward-processing areas such as the anterior cingulate cortex, ventral striatum and insula. To tackle this ongoing debate between homeostatic versus hedonic factors underlying sleep loss-associated weight gain, we rigorously tested the association between SD and food cue processing using high-resolution fMRI and assessment of hormones. After taking blood samples from thirty-two lean, healthy men, they underwent fMRI while performing a neuroeconomic, value-based decision making task with snack food and trinket rewards following a full night of habitual sleep (HS) and a night of SD in a repeated-measures cross-over design. We found that des-acyl ghrelin concentrations were increased after SD compared with HS. Despite similar hunger ratings due to fasting in both conditions, participants were willing to spend more money on food items only after SD. Furthermore, fMRI data paralleled this behavioral finding, revealing a food reward-specific up-regulation of hypothalamic valuation signals and amygdala-hypothalamic coupling after a single night of SD. Behavioral and fMRI results were not significantly correlated with changes in acyl, des-acyl or total ghrelin concentrations. Our results indicate that increased food valuation after sleep loss is due to hedonic rather than hormonal mechanisms.

## Introduction

Numerous epidemiological studies suggest a link between reduced nocturnal sleep and increased risk for overweight and obesity (Patel and Hu, 2008). For example, short sleep duration was positively correlated with body mass index (BMI) in many studies (e.g. Cournot et al., 2004; Heslop et al., 2002; Shigeta et al., 2001; Vioque et al., 2000) and hours of nocturnal sleep were negatively correlated with body fat percentage (Rontoyanni et al., 2007). Besides homeostatic mechanisms, decision-related mechanisms likely contribute to the regulation of food intake (D’Agostino and Small, 2012; Rangel, 2013) and both may consequently affect overeating following sleep loss (Cedernaes et al., 2015; Chaput and St-Onge, 2014).

The dysbalance that sleep loss exerts on homeostasis can be manifested in altered levels of hormones involved in hunger and satiety. Two such candidate hormones are the orexigenic hormone ghrelin and the anorexic hormone leptin. Receptors for ghrelin (Howard et al., 1996) and leptin (Schwartz et al., 1996) are expressed in the hypothalamus, a brain region involved in the regulation of a wide range of biological functions, involving hunger (Anand and Brobeck, 1951) and circadian rhythm (Economo, 1930). For brain activation in response to food stimuli, ghrelin has been found to act as modulator in reward-processing areas (Goldstone et al., 2014; Kroemer et al., 2013; Malik et al., 2008). When also taking sleep loss into account, previous studies found elevated ghrelin concentrations, whereas leptin levels were decreased after sleep restriction (Morselli et al., 2010; Spiegel et al., 2004; Taheri et al., 2004). For example, compared with a night containing a habitual 7-hour sleep period, total ghrelin levels were significantly increased after restricting sleep to 4.5 hours, and even more after a night of total sleep deprivation (SD). The increase in ghrelin was paralleled by increases in subjective hunger (Schmid et al., 2008). However, studies focusing on hormonal changes as an explanation for increased food intake after sleep loss often use very controlled, artificial laboratory environments that do not operationalize realistic life situations, such as feeding with intravenous glucose administrations (Spiegel et al., 2004) or SD while sitting awake in bed the whole night without physical activity (Schmid et al., 2008). Therefore, endocrine modulation has recently been questioned as major contributor to changes in food-related decision-making and the consideration of increased hedonic values of food after SD was brought up as possible explanation for overeating after sleep loss (Chaput and St-Onge, 2014). Thus, control for both homeostatic and hedonic aspects seems crucial in the context of food reward and sleep loss. At the same time, unnecessarily artificial experimental setups might not yield ecologically valid insights.

Functional magnetic resonance imaging (fMRI) studies revealed an increased neural response to food images in regions involved in reward and motivation following total or partial SD, such as the anterior cingulate cortex (Benedict et al., 2012), amygdala (Greer et al., 2013), insula, orbitofrontal cortex, nucleus accumbens and putamen (St-Onge et al., 2012). However, these studies did not examine more general changes in reward processing, e.g. by comparing food rewards to nonfood control rewards, or subjective reward valuation via parametric contrasts. Finally, as discussed in detail above, effects on reward processing might in part be driven by SD-induced hormonal changes (Chaput and St-Onge, 2014; Spiegel et al., 2004). Yet, previous imaging studies lacked control for neuroendocrine factors.

Here we specifically address these issues using combined approaches from decision neuroscience and endocrinology. We recorded fMRI while participants underwent a neuroeconomic decision-making task (BDM auction task) (Becker et al., 1964; Chib et al., 2009) assessing the subjective value of food and nonfood stimuli in a counterbalanced repeated-measure design after habitual sleep (HS) and a single night of total SD. Additionally, we took blood samples before fMRI scanning to determine concentrations of circulating hormones involved in homeostasis. We hypothesized that a full night of SD selectively increases subjective valuation of food vs. nonfood reward, paralleled by selective increases in BOLD signals in reward-and homeostatic-related brain structures. Furthermore, we predicted elevated ghrelin concentrations after SD vs. HS as well as a modulation of behavior and brain activity in response to food stimuli by changes in circulating ghrelin.

## Material and Methods

### Participants

Thirty-two healthy, lean, male participants (mean ± SD age: 26.13 ± 3.80 years, range: 19–33 years; BMI: 23.32 ± 1.44 kg/m^2^, range: 20.52-25.66 kg/m^2^) participated in two fMRI sessions in counterbalanced order, following a single night of SD or HS. No blood samples could be collected from two participants, and the plasma sample of one participant was not frozen immediately after centrifuging in one session, therefore endocrine data are only reported for 29 participants. All participants were right-handed, non-smokers, had normal or corrected-to-normal vision and no history of neurological or psychiatric disorders. On average, they were good sleepers during the last 4 weeks as assed with the Pittsburgh Sleep Quality Index (Buysse et al., 1989) (4.43 ± 0.36). The BMI exclusion criterion was a BMI of 25 kg/m2 or higher. However, if men interested in participating in the study failed this BMI criterion during screening but had at the same time less than 20% body fat, we included these participants in our study. This was the case for 2 participants (BMI: 25.66 and 25.23 kg/m2; percent body fat: 15.3 and 18.4%, respectively). Participants were not on a special diet and did not have any food allergies. All experimental procedures were approved by the local ethics committee (Hamburg Board of Physicians) and all experimental appointments took place at the Institute for Systems Neuroscience at the University Hospital Hamburg-Eppendorf in Hamburg, Germany.

### Overall procedure

Participants visited the institute for three appointments, further described below: one screening and two counterbalanced experimental fMRI sessions with either a HS or a SD condition, separated by one week. The evening before the fMRI scan, they received a standardized dinner at the institute after which they went home to sleep as usual (HS session) or stayed at the institute to spend the whole night awake under constant supervision (SD session).

*Pre-experimental screening session*. After recruitment by online advertisements and a short phone interview, participants were invited to the institute 1–5 days prior to the first experimental session for a pre-experimental screening. During this appointment, participants read the study description carefully and had the opportunity to ask questions concerning the procedure before giving written informed consent. We measured height, weight, body fat percentage, and the familiarity of the stimuli used in the fMRI experiment. Furthermore, a physician conducted a short medical examination to ensure fMRI compatibility. At the end of the screening we did not reveal the order of the two experimental sessions. Additionally, to avoid that participants slept during the afternoon prior to the SD session when HS was the first condition, we instructed them that all combinations of the two conditions could be possible, including two HS or two SD sessions. As a consequence, they did not know if they stayed awake or could go home to sleep until they came in for the evening appointment. After the completion of all three sessions, we uncovered this story.

*Experimental Sessions*. Both experimental sessions started at 8 p.m. with groups of two or three participants undergoing the same experimental condition. Upon arrival, they were told the experimental condition of the night and received a standardized dinner with 741 kcal per serving (pasta with veal strips in cream and mushroom sauce: 582 kcal total, per 100 g serving: 142 kcal, fat: 6.5 g, carbohydrate: 13.2 g, protein 7.7 g; apple: 68 kcal total, per 100g serving: 52 kcal, fat: 0.2 g, carbohydrate: 14.0 g, protein: 0.3 g; strawberry yogurt: 91 kcal total, per 100 g serving: 91 kcal, fat: 3.0 g, carbohydrate: 13.0 g, protein: 2.9 g). Importantly, participants fasted overnight in both conditions, since we instructed them to refrain from food and caloric beverages until the appointment in the morning.

In the HS condition, participants wore an Actiwatch 2 (Philips, Respironics) to track their sleep and wake times until the next morning. They went home to spent the night as usual, and were invited again for the fMRI session the next morning between 7:30 a.m. and 9:30 a.m. In the SD condition, participants stayed at the institute under constant supervision and spent the whole night awake. During this time, they played card games, parlor games, games on game consoles, watched series and movies, and took walks at the university area.

Each experimental session in the morning started with hunger and appetite ratings on a 7-point Likert scale, followed by BDM pre-scan bidding (see section “BDM auction task”), blood drawing immediately before scanning, the BDM choice phase in the scanner and the BDM auction after scanning. We monitored participants in the scanner by online eye-tracking to ensure wakefulness during the fMRI task.

### BDM auction task

Participants performed a Becker-deGroot-Marschak (Becker et al., 1964) auction to assess their willingness to pay (WTP, i.e. subjective value) for a range of snack foods (food reward) and trinkets (nonfood reward) (Chib et al., 2009; Plassmann et al., 2007). In this task, participants have the opportunity to win a trinket and a snack item (factor reward category with levels “food” and “nonfood”). The task consisted of three phases, reported below in more detail: (1) a pre-scan free bidding phase to obtain subjective value estimates for all items, (2) a decision phase in the scanner, and (3) a post-scan auction phase. The procedure for the BDM auction closely followed previous reports (Chib et al., 2009; Plassmann et al., 2007). In particular, care was taken to ensure that all participants understood the auction procedure, and understood that the best strategy is to bid exactly the maximum amount that they are willing to pay for an item in the pre-scan phase. Also, participants were instructed that following scanning on each day, one trial per item category (food and nonfood) from the combined set of trials from the bidding and decision phase would be selected and played out. All snacks and trinkets were available and arranged in the testing room, such that all choices involved the prospect of obtaining the actual rewards.

*Pre-scan bidding phase*. In the bidding phase (see Figure 1 a), participants received 3 € to spend on snacks and 3 € to spend on trinkets. They saw all food and nonfood images and indicated their WTP for each item on a scale from 0 € to 3 € in steps of 0.25 €. They were instructed to bid the maximal amount they were willing to spend on the item and that they could use the full range of the 3 € for each item since only one item per category was drawn in the auction at the end.

**Figure 1.**
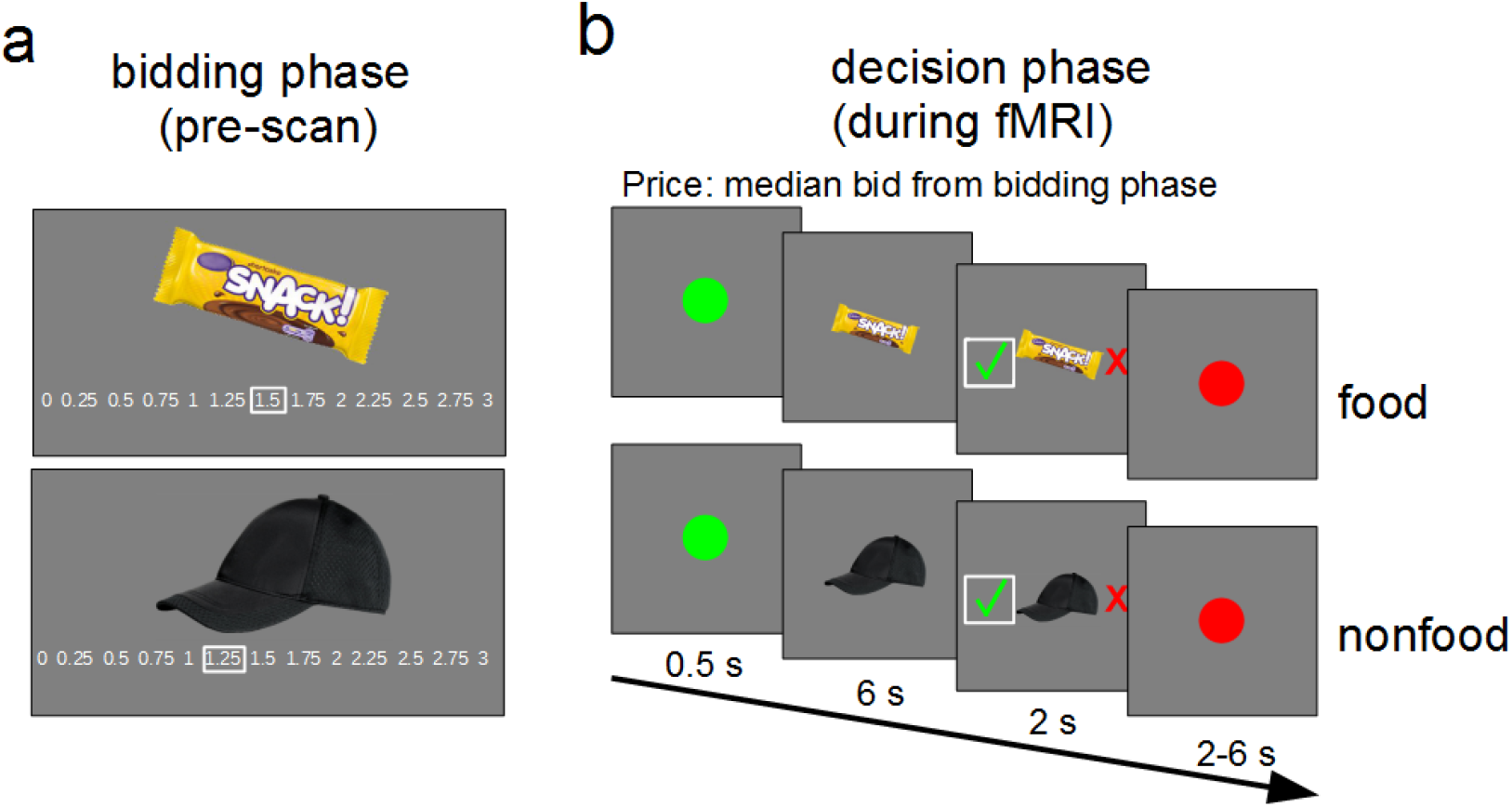
Outline of the BDM task structure of the pre-scan bidding phase (a) and the fMRI decision phase (b).

*fMRI decision phase*. During the choice phase in the fMRI scanner, participants underwent a two-alternative forced choice-task (see Figure 1 b). Participants saw all snack and trinket images again and had to indicate if they would buy or reject them for the reference price, which was the median bid over all food and nonfood items. In a typical trial participants saw a green dot for 0.5 s, followed by the food or nonfood item for 6 s. Subsequently, a red cross (reject) and a green check mark (accept) appeared randomly left and right of the image for 2 s, indicating the decision-making phase. Participants made choices via a MRI compatible button-box. The intertrial interval was marked by a red dot with a randomized presentation time between 2 and 6 s, sampled from a uniform distribution.

*Post-scan auction*. In the auction after scanning one trial per category was randomly drawn from all trials of the bidding and choice phases. The participants’ bid competed against a randomly generated price by the computer between 0 € and 3 € in 0.25 €-steps. If the participants’ price was higher than or equal to that of the computer, they purchased the snack for the lower price and additionally received the difference amount to 3 €. If the computer-generated price was higher than the participants’ price, the item could not be bought but participants received the full 3 €. After the auction, participants stayed another 30 minutes at the institute and could not eat anything except the snack in case they purchased one.

### Stimuli

Visual stimuli consisted of 48 different snack food and 48 different trinket images. The presented snacks were familiar snack foods available in Germany [mean ± SD familiarity: 3.88 ± 0.66, scale from 1 (not familiar) to 5 (highly familiar)], e.g. chocolate bars and chips, taken from an internet search and a previous study (Gluth et al., 2015). Trinkets were familiar [mean ± SD familiarity: 3.64 ± 0.75, scale from 1 (not familiar) to 5 (highly familiar)] everyday items, such as office, drugstore, or university merchandise items, compiled from an internet search and inspired by the trinket items used in Chib et al. (2009).

All images were resized to 400 pixels in the largest dimension, superimposed on a grey background image and presented with Presentation software version 18 (Neurobehavioral Systems, CA, USA). For each participant, half of the 48 stimuli from each category were randomly chosen and presented on the first scanning day, the other half on the second scanning day. Images from both categories were mixed and randomly presented in two runs per session. In the scanner, images were projected on a wall and participants saw them via a mirror mounted on the head coil.

### Behavioral data analyses

*Hunger ratings*. Subjective hunger ratings before the BDM task were compared between sessions with a paired-samples t-test.

*Bids and reaction times*. We assessed differences between bids and reaction times from the BDM pre-scan bidding phase using 2×2 repeated-measures ANOVAs with the factors sleep state (SD, HS) and reward category (food, nonfood).

*Probability to buy*. The probability to buy an item [p(buy)] for the individual median bid during the scanning choice phase was calculated by dividing the number of accepted by the total number of items. Differences between sleep states and reward categories were compared with a repeated-measures ANOVA.

*Hierarchical Bayesian drift diffusion modeling*. To further supplement analyses of bidding behavior, individual participant’s choice and reaction time data were fit with a drift diffusion model (DDM) (Ratcliff and McKoon, 2008). The DDM is a frequently used mathematical model describing two-alternative forced-choice tasks and has been shown to be valid in the context of value-based choices (Mormann et al., 2010). The choice process is modeled as a noisy evidence accumulation process over time between two boundaries. As soon as one of the boundaries is crossed, the associated response is executed. This evidence accumulation is described by several underlying parameters, which in turn give rise to the reaction time distributions for correct and incorrect choices. The precise model formulation can be retrieved from Ratcliff and McKoon (2008). Here, the drift rate *v* reflects the rate of evidence accumulation over time; the boundary separation *a* reflects the amount of evidence required to execute a choice; the non-decision time parameter *t*, captures the time needed to perceive the stimulus and execute a motor response; *z* and corresponds to the starting point of the evidence accumulation process between the two boundaries and was fixed at the midpoint between the two boundaries in the model. In addition, inter-trial variability in non-decision time (*St*) was included in the model, because setting it to zero might be problematic for model estimation (Voss et al., 2015). Correct choices were defined as choices consistent with the median bid from the pre-scan bidding phase, i.e. buying the item if the bid was ≥ median bid and rejection if the bid < median bid, while incorrect choices were defined as choices inconsistent with the median bid from the pre-scan bidding phase.

Parameter values were estimated using a hierarchical Bayesian parameter estimation procedure based on Markov Chain Monte Carlo simulation via the Python-based hierarchical drift diffusion model toolbox (HDDM) (Wiecki et al., 2013). There are two key advantages to this method. First, hierarchical models estimate group and individual parameters simultaneously, allowing the group estimates to constrain the values of individual parameter estimates. Consequently, less trials are needed to reliably estimate individual parameter values. Second, the posterior distributions derived by Bayesian estimation procedures provide the most likely parameter estimates as well as a measure of uncertainty in these estimates.

Two independent but equivalent models were estimated for the HS and the SD conditions. For both models 20,000 samples were drawn from the posterior distribution. 2,000 of these samples were discarded as burn-in. Successful chain convergence was assessed by comparing inter-chain and between-chain variance with the Gelman-Rubin statistic (Gelman, 2004). It was hypothesized that an increase in standardized absolute value difference between the median bid from the bidding phase and the presented stimulus in the choice phase of the task would influence the drift rate differently for the HS compared with the SD condition. This is similar to previous studies that have examined conflict-dependent changes in drift rates associated with SD (Menz et al., 2012). Therefore, drift rates were modeled using a regression approach as a linear function of the standardized absolute value difference in each of the compared models. The linear model used to estimate the trial-by-trial drift rate contained one intercept and a slope parameter for each of the two stimulus types (food vs. nonfood rewards). This HDDM regression approach has been described previously (Wiecki et al., 2013) and has been applied in a number of previous studies (Cavanagh et al., 2011; Herz et al., 2016).

### Blood sampling and analyses

We took blood plasma and serum samples in both sessions to determine circulating levels of ghrelin, leptin, insulin, cortisol, and glucose. Collection of blood samples took place immediately before fMRI scanning, in order to measure endocrine concentrations during the fMRI BDM choice task.

Plasma blood samples of 8.5 ml were collected in BD P800 tubes (BD Biosciences, Heidelberg, Germany) containing K_2_EDTA anticoagulant for ghrelin concentration determination. They were immediately preprocessed by centrifuging at 4 °C for 10 min with 1200 g and pipetting off of the supernatant, and stored at −80 °C.

Serum blood samples of 7.5 ml were collected in Serum Gel Monovettes (Sarstedt, Nümbrecht, Germany) to determine leptin, cortisol, and insulin concentrations. The blood soaked for 45 min in the gel solution before the samples were centrifuged at room temperature for 10 min with 2000 g. The supernatant was pipetted off and stored at −80 °C.

Two additional serum samples of 2.7 ml each for glucose determination were collected in Sarstedt S-Monovettes with Fluorid EDTA (Sarstedt, Nümbrecht, Germany) and also soaked for 45 min in gel solution before centrifuging at room temperature for 10 min with 2000 g. The supernatant was pipetted off and stored at −80 °C.

All serum blood samples were analyzed by the LADR laboratory in Geesthacht, Germany. Leptin was analyzed with a sandwich enzyme-linked immunosorbent assay (ELISA) from DRG (Marburg, Germany), cortisol and insulin with an electro-chemiluminescence immunoassay (ECLIA) method from Roche (Basel, Switzerland), and glucose with a photometric AU 5800 from Beckmann Coulter (Krefeld, Germany).

Total and acyl ghrelin concentrations were analyzed at the Metabolic Core Unit, CBBM, in Luebeck, Germany with a radioimmune assay (Millipore, Billerica, MA, USA).

### Endocrine data analysis

For visualization, raw values of endocrine data were compared using non parametric Wilcoxon signed-rank tests with a two-tailed significance threshold of *p* < 0.05 in Table 1 and Figure 3a, but due to skewed distributions we log-transformed all endocrine parameters prior to further analyses.

**Table 1.** Raw endocrine parameter concentrations after habitual sleep and a full night of SD. Total, acyl, and des-acyl ghrelin as well as serum leptin, cortisol, insulin, and glucose concentrations after a full night of HS or total SD for 29 participants. *Z* values are from non-parametric Wilcoxon signed-rank tests. Values are mean ± s.e.m.

| Endocrine parameter | Habitual sleep<br>(HS) | Sleep deprivation<br>(SD) | z value | p value |
| --- | --- | --- | --- | --- |
| Ghrelin total [pg/ml] | 1048.14 ± 61.54 | 1140.94 ± 64.37 | -1.59 | 0.11 |
| Ghrelin acyl [pg/ml] | 298.60 ± 24.92 | 275.33 ± 23.03 | 0.96 | 0.34 |
| Ghrelin des-acyl [pg/ml] | 749.54 ± 69.06 | 865.61 ± 73.73 | -2.07 | **0.039\*** |
| Leptin [ng/ml] | 1.82 ± 0.21 | 1.77 ± 0.20 | -0.12 | 0.91 |
| Cortisol [mg/l] | 118.00 ± 8.09 | 122.90 ± 7.49 | -0.56 | 0.57 |
| Insulin [μE/l] | 6.53 ± 0.53 | 5.96 ± 0.47 | 0.65 | 0.52 |
| Glucose [mg/dl] | 78.48 ± 2.14 | 79.38 ± 1.86 | -0.02 | 0.98 |

We calculated des-acyl ghrelin concentrations by subtracting raw values of acyl from total ghrelin values.

### fMRI data acquisition

fMRI data were obtained on a Siemens MAGNETOM TRIO 3 T whole-body scanner using a 32-channel head coil. Functional images were collected using single-shot echo-planar imaging with parallel imaging (GRAPPA, in-plane acceleration factor 2) (Griswold et al., 2002) and simultaneous multi-slice acquisitions (“multiband”, slice acceleration factor 2) (Feinberg et al., 2010; Moeller et al., 2010; Xu et al., 2013) as described previously (Setsompop et al., 2012) (TR = 2260 ms, TE = 30 ms, number of slices = 60, flip angle = 80°, voxel size = 1.5 * 1.5 * 1.5 mm). The corresponding image reconstruction algorithm was provided by the University of Minnesota Center for Magnetic Resonance Research.

### fMRI data analysis

*Preprocessing and noise regressors*. Imaging data was analyzed using SPM12 (http://www.fil.ion.ucl.ac.uk/spm/). After removing the first five scans, images were realigned to the first scan and unwarped to account for movement-related effects. The anatomical image scan was then co-registered to the mean realigned EPI image and segmented into grey matter, white matter and CSF. For each subject and session, we additionally computed noise regressors in two steps. First, we created a combined white matter and CSF mask image. Second, we calculated a principal component analysis over all voxels in the mask, identifying all principal components which explained more than 1% of the variance in the mask time series. These principal component scores were included in the first level models as regressors of no interest (mean +/− SEM number of components: SD 8.75 ± 0.3; HS 7.69 ± 0.28).

*First level analyses*. After preprocessing, we set up a general linear model at first level in single-subject space. For each session, we included the following regressors, which were convolved with the hemodynamic response function: 1) onsets of the snack food images as stick function, 2) WTP values on these food items, i.e. subjective values, as first parametric modulator, 3) squared bid values on these food items as second parametric modulator, 4) onsets of the nonfood trinket images as a stick function, 5) WTP values on these nonfood items, i.e. subjective values, as first parametric modulator, 6) squared bid values on these nonfood items as second parametric modulator, 7) regressor of no interest coding for error trials, 8) noise regressors (see above) as additional non-convolved covariates. Normalization parameters were obtained for the structural scans and applied to the single-subject contrast images, writing the normalized contrasts with 1.5* 1.5*1.5mm. Normalized contrast images were smoothed with a 6-mm full-width at half maximum isotropic Gaussian kernel.

*Second level models*. Sleep state-dependent changes in image processing (onset of the stimuli) and valuation signals (parametric effects of WTP) were analyzed on the second level via a flexible factorial model with the within-subject factors reward category (food, nonfood) and sleep state (SD, HS).

*Psychophysiological interaction analysis (PPI)*. We also investigated sleep state x reward category-dependent changes in functional connectivity via PPIs (Friston et al., 1997). For the PPI analysis, we had to bring single subject data into MNI space and therefore re-ran preprocessing as described above with the addition of normalizing the EPIs to MNI space, writing them with 1.5* 1.5*1.5 voxel, and smoothing the images with a 6-mm full-width at half maximum isotropic Gaussian kernel. After preprocessing, we computed first level models and a flexible factorial second level as described above. As seed regions we used peaks in ROIs that reached a FWE-SVC-corrected *p*-value<0.05 in the flexible factorial onset and value interaction contrasts. The peaks are the same as in the initial, single-subject space analysis (right hypothalamus) or very similar (left hypothalamus, right amygdala). The right amygdala [MNI coordinates (x, y, z) = (21, −4, −25)] revealed higher activation for snack onset compared with trinket onset images, and the right (4, −1, −10) and left (−4, −6, −12) hypothalamus clusters showing sleep state x reward category interaction effects with respect to valuation signals.

For the PPI analysis, we proceeded as follows for each subject and each session: We extracted time series in the peak voxels from preprocessed images, created PPI regressors for food onsets and subjective values < nonfood onset and subjective values, and calculated an additional PPI first level model with the user-defined PPI regressors and noise regressors. We than compared activation in response to images and valuation signals between SD and HS groups with a second level paired t-test model.

*Correction for multiple comparisons*. All results are reported using FWE-corrected p-values based on anatomical and/or function regions of interest (ROIs). As ROIs we focused on the hypothalamus, amygdala and vmPFC. A bilateral hypothalamus mask was created using an automated neurosynth metaanalysis with the keyword “hypothalamus” (www.neurosynth.org). We thresholded the resulting reverse inference map at *z* ≥ 18 to exclude unspecific activation and saved it as binary mask image. Note that this approach is not directly based on anatomical considerations, but rather depends on how researchers commonly classify imaging activations as being located in the hypothalamus. Notably, the interaction effect in the right hypothalamus also survived correction across a bilateral anatomical hypothalamus from the WFU Pick Atlas (Maldjian et al., 2003, 2004) [(4, −2, −10): *z*=2.91*, p(FWE)*=0.024] and when coordinate-based ROIs were used as in a previous study (Kullmann et al., 2015) (medial hypothalamus (4, −2, −12) + 2 mm sphere radius; *z*=2.91, *p(FWE)*=0.008). A bilateral amygdala mask was taken from the WFU Pick Atlas, by combining the provided masks for the left and right basolateral amygdalae (Maldjian et al., 2003, 2004). For the vmPFC we used a mask provided by the Rangel Neuroeconomics Laboratory (http://www.rnl.caltech.edu/resources/index.html). This mask is a conjunction of the results of two published metaanalyses (Bartra et al., 2013; Clithero and Rangel, 2014) defining areas that positively correlate with reward value.

For display purposes, activation in fMRI figures is thresholded at *p*<0.005 uncorrected. Activations are projected on a skull-stripped mean anatomical image created with the SPM function *imcalc*. Individual skull-stripped images were created for each participant by using only those voxels of the individual anatomical image matching the segmented white and gray matter images, and after that normalized and averaged.

## Results

### Behavioral Results

Prior to scanning, subjective feelings of hunger did not differ significantly between SD and HS (Figure 2a, *t*_(31)_ =0.75, *p*=0.46). WTP as assessed in the pre-scan auction phase increased selectively for food rewards after SD (sleep state x reward category interaction *F*_(1,31)_=5.48, *p*=0.03, Figure 2b).

**Figure 2.**
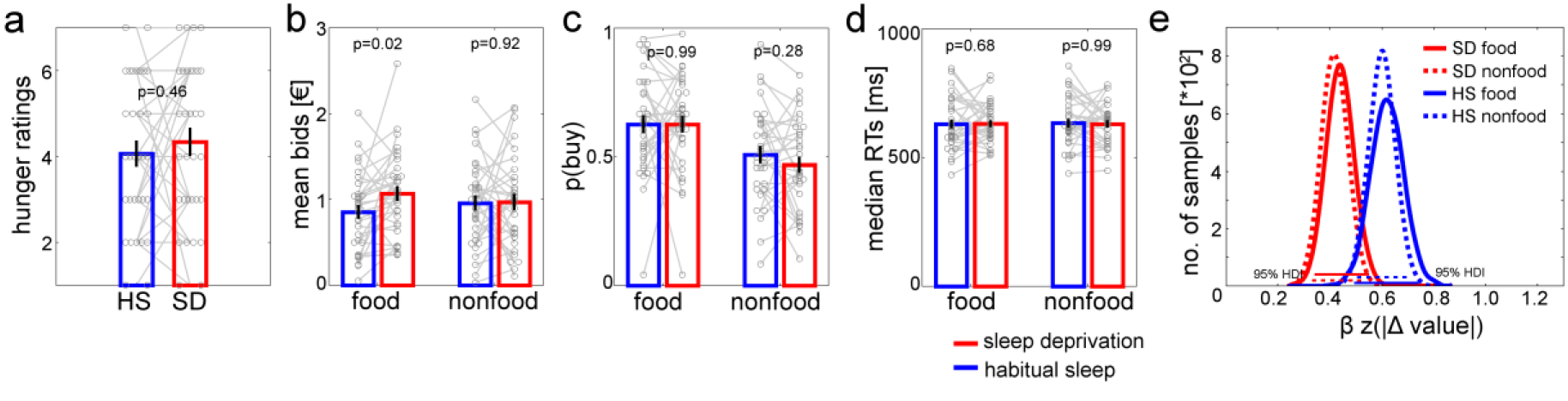
Behavioral and modeling data. Subjective hunger ratings (a) did not differ significantly between HS (blue) and SD (red). After SD, WTP specifically increased for food compared with nonfood (b, sleep state × reward category interaction *p*=0.03). The probability to buy snacks for the reference price during scanning only showed a main effect of category (c, *p*<0.001) and RTs did not differ significantly across sleep and category (d, all *p*>0.58). Diffusion modeling suggested attenuated conflict-induced reductions in drift rate following SD for both food and nonfood rewards (e).

During fMRI, participants made purchasing decisions for all items from the pre-scan bidding phase, with the price set to each participant’s median bid. The probability to buy items at this reference price [(p(buy)] was higher for snacks vs. trinkets (main effect category: *F*_(1,31)_=26.54, *p*<0.001, Figure 2c) but not differentially modulated by sleep (*F*_(1,31)_=0.35, *p*=0.56, Figure 2c) or a sleep × category interaction (*F*_(1,31)_=0.77, *p*=0.39, Figure 2c). There were no effects of sleep state (*F*_(1,31)_=0.05, *p*=0.83, Figure 2d) or reward category (*F*_(1,31)_=0.03, *p*=0.87, Figure 2d) on median reaction times, nor was there an interaction (*F*_(1,31)_=0.6, *p*=0.44, Figure 2d). We further examined choice dynamics using a hierarchical Bayesian implementation (Wiecki et al., 2013) of the drift diffusion model (DDM) (Ratcliff and McKoon, 2008), modeling linear changes in the drift rate *v* across trials *t* as a function of |*WTP(t)—medianbid*| using a full DDM with a linear term for *v*. Descriptively, conflict-dependent slowing tended to be more pronounced for SD compared to HS (see Figure 2e, but note that the 95% highest density intervals for the regression coefficients showed some overlap). This is in line with previous findings (Menz et al., 2012) and suggests that valuation (i.e. bidding behavior) but not choice dynamics (drift rate) were modulated in a reward-category specific fashion following SD.

### fMRI Results

Functional MRI revealed a main effect of subjective value across sleep states and reward categories in ventromedial prefrontal cortex (vmPFC; MNI coordinates (x, y, z)=(−9, 41, −7); *z*=4.78, *p(FWE)*=0.001; see Figure 3). In contrast, bilateral hypothalamus and amygdala both showed distinct sleep x reward category interaction effects: Valuation signals (i.e. parametric effects of WTP on BOLD amplitude) in bilateral hypothalamus (right (4, −1, −10): *z*=3.27, *p(FWE)*=0.018; left (−4, −7, −12): *z*=2.92, *p(FWE)*=0.048) were selectively increased for food rewards following SD (Figure 4). Additionally, stimulus onset-related activity in bilateral amygdala (right (20, −2, −25): *z*=3.48, *p(FWE)*=0.04; left (−24, −7, −25): *z*=3.19, *p(FWE)*=0.088, see Figure 5) was selectively enhanced for food rewards following SD. Psychophysiological interaction analysis (PPI) then revealed that these same regions also showed a food-specific increase in functional connectivity following SD vs. HS (coupling of right amygdala with left hypothalamus (−6, −7, −12): *z*=3.50, *p(FWE)*=0.011; with right hypothalamus (9, −4, −12): *z*=2.67, *p(FWE)*=0.098, see Figure 6).

**Figure 3.**
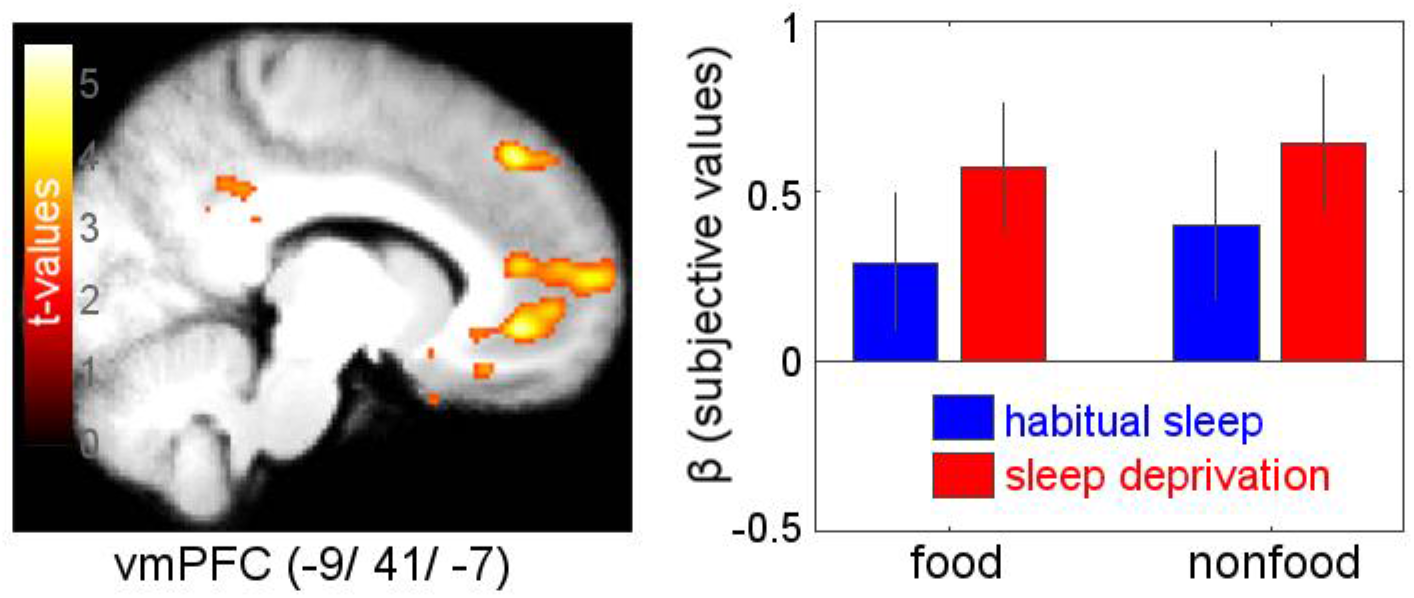
Main effect of subjective reward value (WTP) across reward categories (food and nonfood) and states (HS, SD) revealed effects in ventromedial prefrontal cortex [(−9, 41, −7): *z*=4.78, *p(FWE)*=0.001].

**Figure 4.**
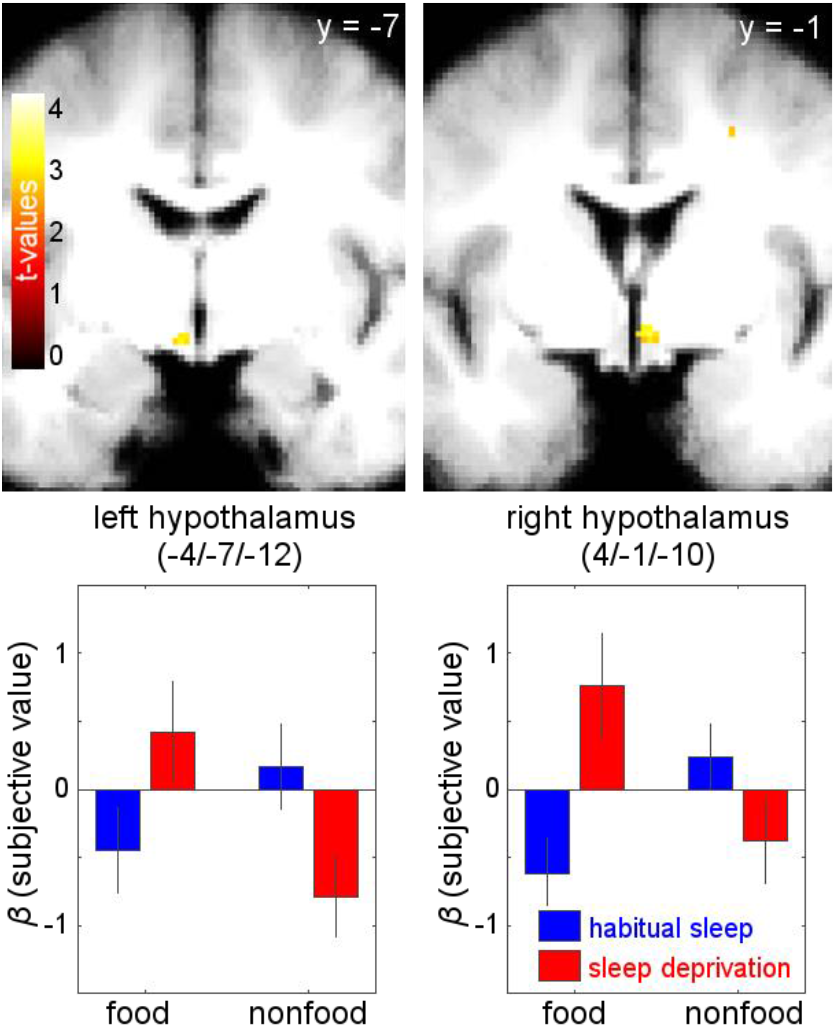
Parametric value effects on BOLD amplitude. Sleep state x reward category interaction contrast for reward valuation signals revealed food-specific valuation increases in bilateral hypothalamus following SD (right (4, −1, 10): *z*=3.27, *p(FWE)*=0.018; left (−4, −7, −12): *z*=2.92, *p(FEW)*=0.048; display threshold: *p*<0.005, uncorrected).

**Figure 5.**
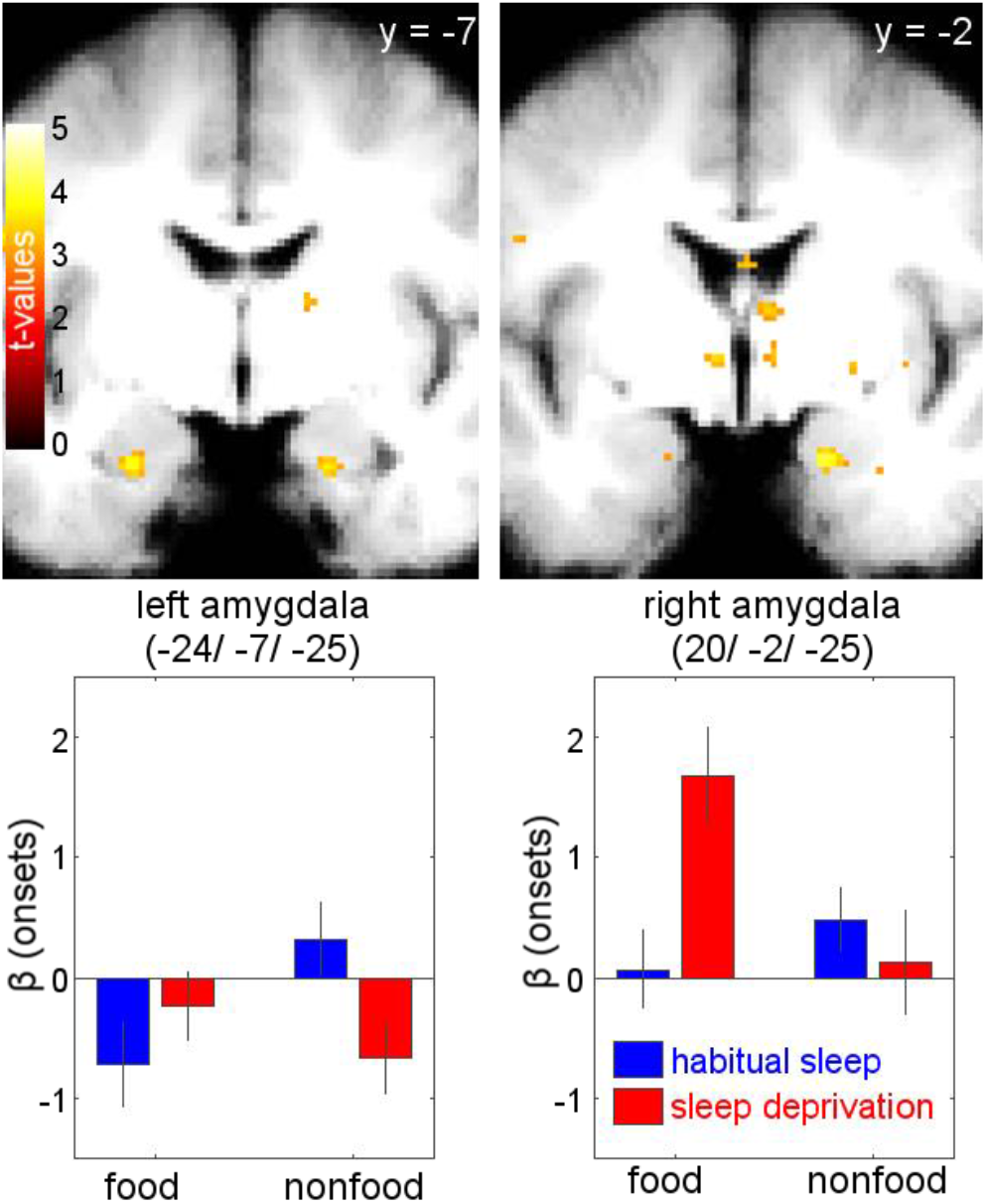
Image onset effects on BOLD amplitude. Sleep state x reward category interaction with respect to activation related to image presentation revealed increased activation in bilateral basolateral amygdala for food vs. nonfood rewards following SD vs. HS (right (20, −2, −25): *z*=3.48, *p(FWE)*=0.04; left (−24, −7, −25): *z*=3.19, *p(FWE)*=0.088; display threshold: *p*<0.005, uncorrected).

**Figure 6.**
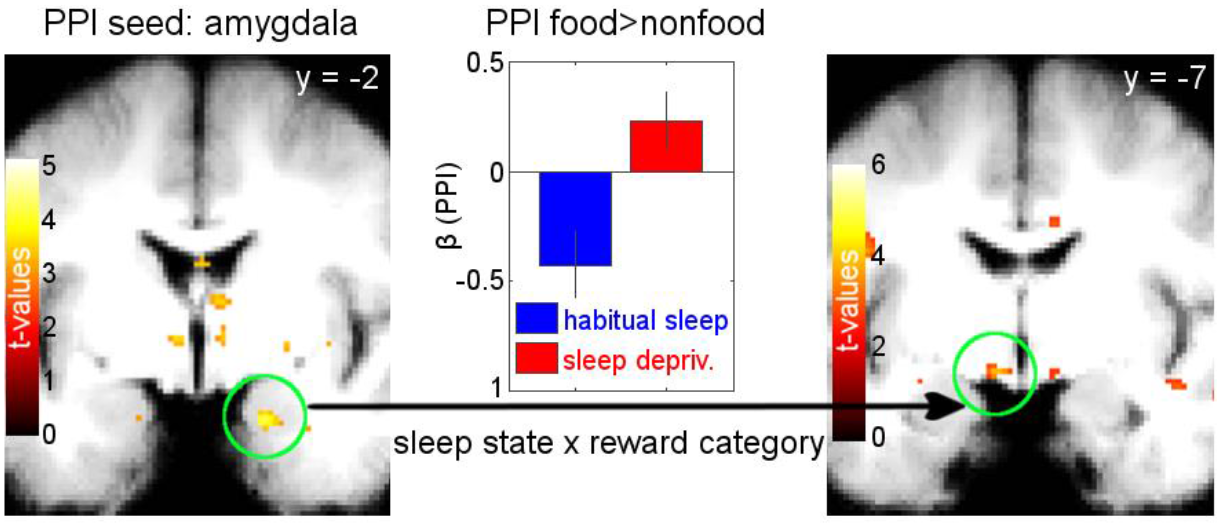
Functional connectivity results from a psycho-physiological interaction analysis (PPI). The seed was placed in the right amygdala peak from the onset contrast analysis, showing higher activation for food vs. nonfood items following SD vs. HS (left panel). Coupling of this seed with left hypothalamus (left (−6, −7, −12): *z*=3.50, *p(FWE)*=0.011; right panel) was significantly greater for food vs. nonfood in SD vs. HS. The center panel shows the food vs. nonfood coupling parameters for SD and HS.

Tables of all activated brain regions in each of the four above described contrasts are appended at the end of the manuscript (Supplemental Tables 1–4).

### Endocrine modulation

Blood samples collected immediately before fMRI scanning revealed that SD induced a distinct morning increase in des-acyl ghrelin (Wilcoxon signed-rank test: *z*=2.27, *p*=0.04, see Figure 7a), while morning levels of total ghrelin, acyl ghrelin, leptin, insulin, glucose and cortisol did not differ significantly between SD and HS (all *p*>=0.11, see Table 1). Furthermore, des-acyl ghrelin concentrations of SD and HS were significantly correlated within participants (*R*=0.71, *p*<0.001, see Figure 7b).

**Figure 7.**
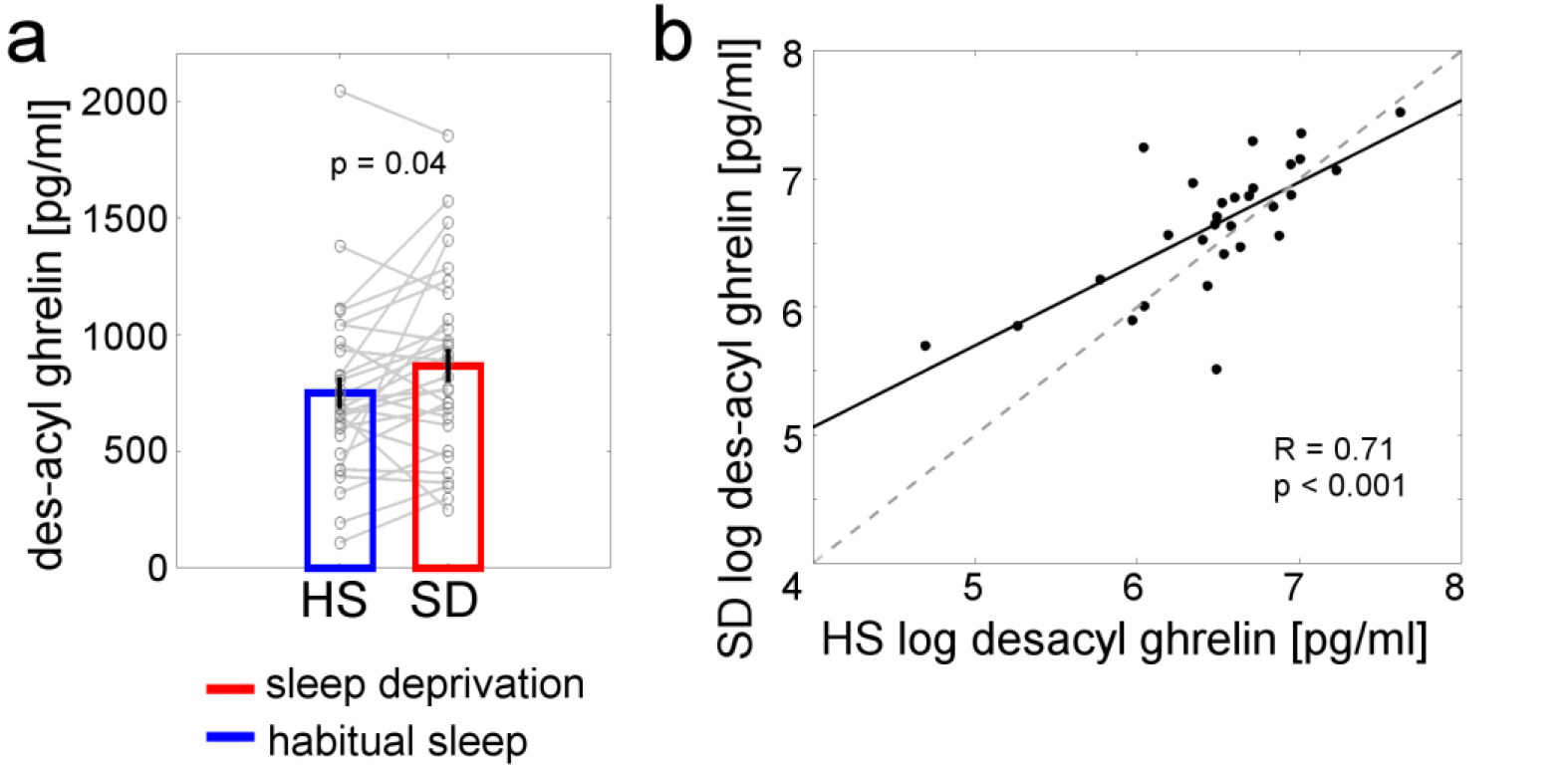
Des-acyl ghrelin data. (a) Blood plasma concentrations of des-acyl ghrelin raw values were higher after SD compared with HS (*z*=2.27, *p*=0.04). (b) Log-transformed des-acyl ghrelin plasma concentrations were significantly correlated within participants between the SD and the HS conditions (*R*=0.71, *p*<0.001).

Since ghrelin was the only endocrine parameter that differed between the two conditions, only the three ghrelin parameters were considered in further analyses. The magnitude of the intra-individual increases in ghrelin after SD vs. HS did neither correlate with the change in WTP for food items (all |*R*|<0.04, all *p*>0.87), nor with the change of the difference between WTP for food and nonfood items (all |*R*|<0.31, all *p*>0.11). For hunger ratings, the change from HS to SD did neither correlate with the change in total nor in acyl ghrelin (all |*R*|<0.26, all *p*>0.18). However, the change in des-acyl ghrelin from HS to SD revealed a significant negative correlation with the change in hunger ratings (*R*=–0.42, *p*=0.02). This correlation did not withstand correction for multiple comparisons, i.e. taking into account the analysis of three different behavioral measures. When regarding only the single sessions, there was no significant correlation between hunger ratings and des-acyl ghrelin values in the SD session (*R*=0.12, *p*=0.53), but in the HS session (*R*=−0.48, *p*=0.009).

Concerning fMRI effects, the change in ghrelin between SD and HS sessions did neither correlate with valuation effects in hypothalamus (all |*R*|<0.25, all *p*>0.18) nor with onset-related activity in amygdala (all |*R*|<0.34, all *p*>0.07), nor with the coupling between right amygdala and left hypothalamus (all |*R*|<0.08, all *p*>0.68).

## Discussion

We found that a full night of SD compared with a night of HS increased the subjective values of snack food rewards compared with nonfood rewards. This behavioral result was paralleled by increased amygdala and hypothalamus activity selectively after SD in response to food image onsets or their parametrically modulation by value. Furthermore, the connectivity between amygdala and hypothalamus was increased after SD for food rewards. However, we could not replicate hormonal modulations of brain activity in response to food after SD.

In this study, we used a well-established paradigm from behavioral economics, the BDM auction task (Becker et al., 1964). This task has been successfully used in conjunction with food and nonfood rewards (Chib et al., 2009; Plassmann et al., 2007) revealing vmPFC as region of common representation of WTP for rewards from different categories (Chib et al., 2009). We replicated this result and observed value-related vmPFC activation in essentially the same location as a previous study [peak coordinates (x, y, z), main effect of value present study: (−9, 41, −7); Chib et al., 2009: (−9, 39, −6)].

The results of our study complement another experiment using a parametric design to examine the effect of SD on food valuation. Greer et al. (2013) reported parametric modulations of food image reactivity by WTP in the amygdala, whereas we found this area involved in processing food images after SD without parametric modulation. However, hypothalamus activity was parametrically modulated after SD in response to food images. Importantly, the nonfood reward control condition allowed us to account for general differences in brain reward responsivity between the two states.

None of the previous studies investigating SD in combination with food image stimuli did control for endocrine effects on BOLD activity (Benedict et al., 2012; Greer et al., 2013; St-Onge et al., 2012), despite the association that several previous studies found between ghrelin and neural food reward responses (Goldstone et al., 2014; Kroemer et al., 2013; Malik et al., 2008). As expected, we observed increases in circulating ghrelin for SD compared with HS. However, only des-acyl ghrelin increased after SD, whereas total and acyl ghrelin levels were not significantly different between conditions. The des-acylated form of ghrelin was initially thought to be an inactive degradation product of acyl ghrelin (Kojima et al., 1999). Although this view has recently been questioned (Delhanty et al., 2012, 2014; Müller et al., 2015), the exact mechanisms of action of des-acyl ghrelin still need to be elucidated. The interpretation of our results is complicated by the inconsistent use of the ghrelin parameters in previous studies, some of which exclusively focused on acyl ghrelin (Goldstone et al., 2014) or des-acyl ghrelin (Kroemer et al., 2013), or indirectly determined ghrelin levels via growth hormone assessment (Malik et al., 2008). Thus, future studies should try to include all ghrelin parameters if possible or carefully explain why they focus on a certain form of ghrelin.

Contrary to our hypothesis, changes from SD to HS sessions in the three ghrelin parameters were not correlated with changes in fMRI activity. For des-acyl ghrelin, changes even negatively correlated with changes in hunger ratings. The unexpected negative correlation with the change in hunger ratings was driven by the HS session and did not withstand correction for multiple testing. In addition to the absence of associations between ghrelin level changes and other behavioral data changes as well as between changes in fMRI activity, this result further argues against a hormonal account of our behavioral findings. Regarding functional imaging data, no correlations with changes in ghrelin levels could be found. One potential explanation might be a ceiling effect of homeostatic hunger in both conditions. Average total ghrelin levels were >1000 pg/ml in both HS and SD, which is substantially higher than in previous studies (e.g. 850 pg/ml after total SD and 720 pg/ml after 7 hours of sleep in Schmid et al., 2008). Similarly, for des-acyl ghrelin average blood plasma concentrations of 760 pg/ml were higher than in other overnight fasting studies (e.g. 410 pg/ml in Kroemer et al. 2013 for male participants). The lacking homeostatic effects on brain activation after SD are in line with previous work demonstrating the absence of ghrelin modulation of amygdala-hypothalamus interactions, which likely reflect an increased hedonic versus homeostatic drive (Sun et al., 2015). Thus, it seems reasonable to conclude that hedonic factors might additionally contribute to increased food valuation after sleep loss. Another potential modulating parameter involved in selectively increased food valuation after SD could be dopamine (Volkow et al., 2012), and future studies might further explore the potential contribution of alterations in dopamine transmission using PET imaging.

Our analyses of both choice and response time data using DDM and a regression approach revealed attenuated value-dependent changes in the drift rate following SD. This observation is consistent with previous findings (Menz et al., 2012; Ratcliff and Van Dongen, 2009, 2011). However, DDM parameters did not show differential effects for food vs. nonfood rewards, suggesting that choice dynamics were affected by SD but not in a category-specific fashion. We also note that our design was not optimized for the application of the DDM, since for example we imposed a minimum waiting period prior to response logging. Nonetheless, we observed effects of SD similar to previous studies (Menz et al., 2012; Ratcliff and Van Dongen, 2009, 2011).

One problem of modern society is sleep restriction building up constantly over a week of work (McCoy and Strecker, 2011) rather than total SD. However, in our approach we used a total night of using SD is a first step in unravelling homeostatic and/or hedonic mechanisms underlying sleep loss-associated increased food valuation. Potential differences in the effects of partial and total SD on homeostasis-related processes should be directly explored in future studies.

A limitation of our study is the lack of reliable objective measures for sleep quality and quantity. In the present study we used SD as intervention to operationalize sleep loss and consequently focused on the effects of SD on food valuation. Therefore, participants remained under constant supervision throughout the SD night to ensure compliance with all protocols. Since our main research question did not aim at investigating possible effects of the active role of sleep in food valuation, HS control sleep was not monitored in detail (e.g. via polysomnography) and only actigraphy was applied to monitor sleep duration before HS sessions, in combination with a questionnaire-based assessment of general sleep quality during the last month. Future studies might use more reliable objective sleep measures such as polysomnography to control sleep quality and quantity, and to explore a potential active role of sleep quality in modulating homeostasis-related processes.

Taken together, our findings reveal a mechanism through which SD might promote food intake by enabling food cues to gain access to processing in hypothalamic circuits via the amygdala. This might then drive a food-specific increase in behavioral valuation as well as hypothalamic representations of value, thereby potentially increasing the likelihood of overeating and consequentially weight gain and obesity risk.

## Author contributions

J.P., J.S.R. and L.S. designed research, J.S.R. and M.M.M. performed research, J.S.R., L.B. and J.P. analyzed data, H.S. and S.M.S. contributed analytical tools, J.P. and J.S.R. drafted the paper and all authors provided revisions.

## Acknowledgements

We thank Sophie Klusen for help with data acquisition.

---

**Supplemental Table 1:** Brain regions where activity increased as a response of subjective value for both categories (food and nonfood) and both states (SD and HS), for *p* < 0.001 with at least 5 contiguous voxels.

| Region | MNI Coordinates (x y z) | z value |
| --- | --- | --- |
| L ventromedial prefrontal cortex/ medial orbitofrontal frontal cortex | -9 41 -7 | 4.78 |
| L angular gyrus | -50 -62 40 | 4.47 |
| L posterior cingulate gyrus | -3 -50 20 | 4.24 |
| R inferior occipital gyrus | 42 -84 -8 | 3.87 |
| R inferior frontal gyrus, par orbitalis | 28 26 -8 | 3.78 |
| L inferior frontal gyrus, par orbitalis | -46 26 -8 | 3.68 |
| R caudate nucleus | 10 11 16 | 3.61 |
| L gyrus rectus | -10 30 -19 | 3.53 |
| L middle frontal gyrus | -34 50 -8 | 3.49 |
| L inferior temporal gyrus | -52 -28 -20 | 3.49 |
| R middle cingulate gyrus | 3 -36 32 | 3.48 |
| L angular gyrus | -38 -66 32 | 3.48 |
| L inferior frontal gyrus, pars triangularis | -38 23 2 | 3.48 |
| L insula | -28 16 -8 | 3.45 |
| L angular gyrus | -46 -50 23 | 3.42 |
| L caudate nucleus | 0 8 16 | 3.40 |
| R anterior cingulate gyrus | 8 28 -7 | 3.40 |
| L superior frontal gyrus | -21 52 -12 | 3.40 |
| R insula | 30 17 -19 | 3.36 |
| R middle frontal gyrus | 44 40 11 | 3.35 |
| L middle temporal gyrus | -68 -32 -2 | 3.34 |
| R inferior temporal gyrus | 50 -1 -34 | 3.27 |
| L middle cingulate gyrus | -8 -43 34 | 3.27 |
| L middle temporal gyrus | -63 -34 -16 | 3.23 |

**Supplemental Table 2:** Brain regions where activity selectively increased for food compared with nonfood reward values after SD compared with HS, for *p* < 0.001 with at least 5 contiguous voxels.

| Region | MNI Coordinates (x y z) | z value |
| --- | --- | --- |
| L parahippocampal gyrus | -10 -16 -21 | 3.89 |
| R gyrus rectus | 16 28 -12 | 3.46 |
| L precuneus | -22 -49 12 | 3.42 |
| R middle frontal gyrus, orbital part | 16 42 -13 | 3.35 |
| R middle frontal gyrus | 27 4 41 | 3.35 |
| R hypothalamus | 4 -1 -10 | 3.27 |

**Supplemental Table 3:** Brain regions where activity selectively increased with the onset of food compared with nonfood images after SD compared with HS, for *p* < 0.001 with at least 5 contiguous voxels.

| Region | MNI Coordinates (x y z) | z value |
| --- | --- | --- |
| L middle temporal gyrus | -36 6 -30 | 4.19 |
| L gyrus rectus | -16 23 -12 | 4.12 |
| L posterior cingulate gyrus | -10 -44 24 | 3.99 |
| L precuneus | -6 -62 46 | 3.89 |
| R superior temporal pole | 34 11 -28 | 3.80 |
| R middle temporal gyrus | 66 -10 -24 | 3.77 |
| L superior temporal gyrus | -36 -26 -1 | 3.77 |
| L hippocampus | -30 -28 -8 | 3.75 |
| R anterior cingulate gyrus | 9 24 16 | 3.65 |
| L posterior cingulate gyrus | -9 -32 20 | 3.63 |
| L precuneus | -15 -43 4 | 3.61 |
| L middle frontal gyrus | -34 42 17 | 3.60 |
| L anterior cingulate gyrus | -8 30 14 | 3.58 |
| L parahippocampal gyrus | -6 -22 -25 | 3.53 |
| L anterior cingulate gyrus | -4 42 17 | 3.51 |
| L middle cingulate gyrus | -6 -42 40 | 3.49 |
| L posterior cingulate gyrus | -9 -34 30 | 3.49 |
| R amygdala | 20 -2 -25 | 3.48 |
| L middle temporal gyrus | -52 -40 -10 | 3.45 |
| R thalamus | 6 -13 -2 | 3.39 |
| R hippocampus | 20 -37 0 | 3.39 |
| L hippocampus | -26 -7 -26 | 3.29 |
| L postcentral gyrus | -52 -8 24 | 3.21 |
| R angular gyrus | 42 -60 41 | 3.19 |

**Supplemental Table 4:** Brain regions that were coupled with right amygdala and that increased with the onsets of food compared with nonfood images after SD compared with HS, for *p* < 0.001 with at least 5 contiguous voxels.

| Region | MNI Coordinates (x y z) | z value |
| --- | --- | --- |
| L fusiform gyrus | -24 -36 -22 | 4.89 |
| L posterior orbitofrontal cortex | -26 29 -16 | 4.81 |
| L middle frontal gyrus | -44 28 30 | 4.25 |
| L thalamus | -14 -16 -8 | 3.97 |
| L precuneus | -10 -50 12 | 3.85 |
| R thalamus | 6 -12 -12 | 3.84 |
| L anterior cingulate gyrus | -8 35 28 | 3.81 |
| R insula | 26 16 -14 | 3.75 |
| L middle cingulate gyrus | -3 -13 30 | 3.73 |
| R angular gyrus | 34 -42 35 | 3.69 |
| R parahippocampal gyrus | 18 -36 -6 | 3.68 |
| L hypothalamus | -8 -8 -12 | 3.67 |
| R precuneus | 22 -54 34 | 3.65 |
| R insula | 36 10 -13 | 3.52 |
| R middle temporal gyrus | 39 -66 14 | 3.49 |
| L lobule IV, V of cerebellar hemisphere | -9 -52 -24 | 3.48 |
| L hippocampus | -34 -16 -12 | 3.47 |
| R posterior orbitofrontal cortex | 26 24 -16 | 3.46 |
| L middle occipital gyrus | -46 -78 17 | 3.43 |
| L middle frontal gyrus | -26 40 28 | 3.40 |
| L parahippocampal gyrus | -27 -16 -24 | 3.38 |
| R middle cingulate gyrus | 3 -38 34 | 3.37 |
| L superior frontal gyrus | -20 12 36 | 3.32 |
| L middle cingulate gyrus | -8 -34 34 | 3.24 |
| R middle temporal gyrus | 44 -66 20 | 3.21 |
| L middle temporal gyrus | -46 -38 -6 | 3.18 |

